# Acute caffeine ingestion differentially alters motor unit behaviour during submaximal and fatiguing contractions

**DOI:** 10.64898/2026.09.15.751700

**Authors:** Yuxiao Guo, Thomas B. Inns, Vic Sun, Eleanor J. Jones, Bethan E. Phillips, Philip J. Atherton, Mathew Piasecki

## Abstract

**Background:** Caffeine is widely used as an ergogenic aid, yet its effect on neuromuscular function remains heterogeneous across exercise conditions. Although caffeine is known to enhance central nervous system excitability through adenosine receptor antagonism, the extent to which these effects modify spinal motor output remains unclear. Therefore, the present study aimed to characterise acute caffeine-induced alterations in neuromuscular properties during submaximal and sustained fatiguing contractions of the knee extensors.

**Methods:** Seventeen healthy adults completed a randomised, placebo-controlled crossover study (3 mg/kg caffeine). Circulating serum caffeine concentrations were periodically quantified, while motor unit (MU) discharge behaviour, and ΔF were derived from high-density surface electromyography. Statistical significance was set at p = 0.05.

**Results:** No intervention effects were observed for MVC, force steadiness, or time to task failure (all p > 0.05). During graded contractions, vastus lateralis MU discharge rate was higher under caffeine (p < 0.001), and discharge variability was lower (p = 0.016). Recruitment and derecruitment thresholds were significantly greater under caffeine at 50% MVC (both p < 0.012), whereas paired MU analyses of ΔF was unchanged (p = 0.083). During sustained contractions, caffeine altered fatigue-related trajectories of both MU discharge and discharge variability (intervention × phase interaction, p < 0.011).

**Conclusions:** Acute caffeine ingestion alters motor unit behaviour without enhancing maximal force or fatigue resistance. These effects vary with contraction intensity and fatigue state but are not accompanied by a detectable change in discharge rate hysteresis in the vastus lateralis.

## Introduction

Caffeine is one of the most widely used ergogenic aids and has consistently been shown to improve aspects of exercise performance, including reduced perceived exertion, enhanced force production, and prolonged exercise tolerance (Grgic et al., 2018; Guest et al., 2021). However, these performance benefits are not uniformly expressed, with substantial variability observed between individuals, muscle groups and exercise tasks (Pickering & Grgic, 2019). Although the behavioural effects of caffeine are well established, they provide limited insight into the neural processes responsible for these task-dependent responses.

Caffeine exerts its primary central action through antagonism of adenosine A_1_ and A_2_A receptors (Fredholm et al., 1999; McLellan et al., 2016), thereby reducing adenosine-mediated inhibition and increasing cortical excitability. Consistent with this mechanism, neurophysiological studies have demonstrated enhanced corticospinal excitability following caffeine ingestion (Specterman et al., 2005; Strunge et al., 2024). However, increased corticospinal excitability does not consistently translate into improved motor performance (Guest et al., 2021), suggesting that additional neural processes downstream of the cortex contribute to the heterogeneous ergogenic responses observed across exercise tasks. As descending cortical commands must ultimately be integrated and transformed within the spinal motoneuron pool before force can be generated, the motoneuron pool represents the critical physiological interface through which supraspinal excitation is converted into functional motor output.

Volitional force is generated through the recruitment and discharge of motor units (MUs), reflecting the transformation of descending and peripheral inputs by the motoneuron pool (Farina & Negro, 2015). MU firing is further shaped by intrinsic motoneuron properties, particularly persistent inward currents (PICs), which amplify and prolong depolarisation and are strongly regulated by monoaminergic neuromodulation, including serotonergic and noradrenergic pathways (Heckman et al., 2008). Accordingly, concurrent assessment of MU discharge behaviour and ΔF via paired MU analysis (Mesquita et al., 2024) provides a means of determining whether caffeine-related changes in motor output are accompanied by measurable estimates of PIC-related intrinsic motoneuron excitability.

Consistent with this mechanistic framework, previous studies have investigated neural mechanisms underlying caffeine-induced changes in motor performance. However, evidence at the level of MU output and intrinsic motoneuron excitability remains incomplete and inconsistent. Caffeine did not alter ΔF in the tibialis anterior across different contraction intensities or following repeated fatiguing contractions (Mackay et al., 2023), whereas more recent evidence suggests muscle-specific effects on motoneuron output, with caffeine enhancing responses in the soleus but not the medial gastrocnemius (Popesco et al., 2026). Since PICs are strongly regulated by monoaminergic inputs, including serotonergic modulation (Heckman et al., 2008; Rekling et al., 2000), circulating serotonin concentrations may provide additional context for interpreting potential changes in motoneuron excitability (Perrier et al., 2013). Evidence regarding MU discharge behaviour is also mixed, with studies reporting variable effects of caffeine on motor unit firing properties across different experimental conditions (Ghazaleh et al., 2024; Lin et al., 2025). Collectively, whether caffeine-induced changes in VL MU output are accompanied by changes in intrinsic motoneuron excitability remains unclear.

Therefore, the present study aimed to characterise the effects of acute caffeine ingestion on MU behaviour and intrinsic motoneuron excitability during graded submaximal and sustained fatiguing contractions. Specifically, we examined MU discharge rate and variability, recruitment and derecruitment behaviour, and peak discharge rate, together with ΔF as an index of PIC-related intrinsic motoneuron excitability.

## Methods

### Participants and ethics approval

Seventeen healthy adults (13 males, 4 females; mean age = 22.59 ± 4.37 years) were recruited from the local community through advertisements. All participants provided written informed consent prior to participation.

Participants underwent clinical screening prior to enrolment to exclude conditions that could affect neuromuscular function or participant safety. Exclusion criteria included metabolic disease, lower limb musculoskeletal disorders, acute cardiovascular or cerebrovascular disease, active malignancy, uncontrolled hypertension, and the use of medications known to influence neuromuscular or vascular function.

This study was approved by the University of Nottingham Faculty of Medicine and Health Science Research Ethics Committee (188-0221) and was conducted in 2022-23 in accordance with the Declaration of Helsinki.

### Experimental design and protocol

This study employed a single-blind, randomised, cross-over study design. Following preliminary screening to confirm eligibility, all participants completed two distinct experimental sessions, with a 7-day washout interval implemented between trials. To mitigate potential confounding variables that may skew neuromuscular and blood biomarker outcomes, participants were instructed to refrain from strenuous exercise for 72 hours prior to each testing session and completely abstain from alcohol and caffeine-containing products for 24 hours preceding each trial.

Prior to the intervention procedures, baseline anthropometric and muscle morphological assessments were completed. Body mass and standing height were measured using calibrated scales and a stadiometer, respectively, and body mass index (BMI) was calculated as body mass divided by height squared. Vastus lateralis (VL) cross-sectional area (CSA) was assessed using B-mode ultrasonography at the anatomical midpoint of the muscle, as previously described (Guo et al., 2022). Three transverse images were analysed using ImageJ software (National Institutes of Health, USA), and the mean value was used to represent VL CSA.

Each testing day began at 0900h. A trained clinician performed peripheral venous cannulation to facilitate repeated blood sampling throughout the experimental session. A baseline blood sample (5 ml) was collected immediately following cannulation. Participants then consumed the assigned 500 ml test beverage within 5 min under direct supervision. The test beverage consisted of 450 ml purified water and 50ml orange squash; for the caffeine intervention condition, anhydrous caffeine powder (3 mg/kg body mass) was added to the base solution, while the placebo condition contained no active ingredients. The orange squash was incorporated to mask the distinct bitter taste of caffeine, ensuring the two test beverages were matched for taste and appearance and indistinguishable by sensory cues alone.

Following beverage ingestion, several venous blood samples were collected at□approximately 30 min and 50 min post-ingestion. During this post-ingestion period, participants completed a standardised bout of neuromuscular assessments, with detailed protocols described below.

### Experimental procedures

#### Force recordings

Participants were seated in a custom-built chair with hips and knees flexed at approximately 90°. The lower leg was securely fastened to a calibrated force dynamometer via non-elastic strapping positioned proximal to the medial malleolus, eliminating extraneous limb movement. A pelvic seat belt was fastened to stabilize the upper trunk during maximal and submaximal contractions.

Following a standardised warm-up involving several submaximal isometric contractions, participants performed maximal isometric voluntary contractions (MVCs). To prevent compensatory movements, participants maintained their arms crossed over the chest and were instructed not to grip the chair sides throughout testing. Real-time visual feedback of force output was displayed on a front-facing monitor, and standardised verbal encouragement was delivered to encourage maximal voluntary effort.

Participants completed 2–3 additional MVC attempts with a 60s rest interval between trials to ensure sufficient recovery. Trials were considered valid if the peak force difference between the final two consecutive attempts was less than 5%; the highest peak force value (in Newtons) across valid trials was defined as the participant’s baseline MVC and used for subsequent test normalisation.

### Neuromuscular protocols

#### Submaximal trapezoidal contractions

Following baseline MVC assessment, participants performed a series of submaximal trapezoidal isometric contractions at 10%, 25%, 40%, and 50% MVC. Each contraction comprised a 5-s ramp-up, a 12-s constant plateau at the target intensity, and a 5-s ramp-down. Participants additionally performed two triangular ramp contractions to 20% MVC, each consisting of a 10-s linear ramp-up followed immediately by a 10-s linear ramp-down. The trial with smoother force profile was selected for subsequent analysis.

#### Sustained fatiguing contraction

Participants were instructed to sustain a target force corresponding to 30% MVC for as long as possible. Task failure was defined as the point at which force output fell and remained 10% below the target intensity for 3 consecutive seconds. Immediately following the fatiguing contraction (within ∼5s), an additional MVC was performed and defined as the post-task MVC. Baseline and post-task MVC values were subsequently compared to assess the change in maximal voluntary force across the experimental protocol.

### High-density surface electromyography (HDsEMG)

A semi-disposable HDsEMG electrode array (64 electrodes, 13□×□5, 8 mm, I.E.D., GR08MM1305, OT Bioelettronica, Inc., Turin, Italy) was positioned over the muscle belly of the right vastus lateralis. The array was aligned parallel to the proximal-to-distal orientation of the underlying muscle fascicles and secured to the skin using flexible tape. The adhesive grids were attached to the surface of the muscle by disposable bi-adhesive foam layers (SpesMedica, Bettipaglia, Italy). The skin electrode contact was facilitated by filling the cavities of the adhesive layers with conductive paste (AC Cream, SpesMedica). A strap ground electrode (WS2, OTBioelettronica, Turin, Italy) dampened with water was positioned around the ankle of the right leg to optimise the signal quality. To ensure electrode positioning reproducibility across visits in this cross-over design, the distances from the four corners of the electrode array to the knee joint were precisely measured and marked. These reference markers were used to ensure consistent electrode placement in subsequent trials.

HDsEMG signals were recorded in a monopolar configuration, amplified (×□256), band-pass filtered at 10-500 Hz, and digitized at 2000 Hz using a 16-bit wireless amplifier (Sessantaquattro, OTBioelettronica, Turin, Italy). Raw signals were exported from the OTBioLab software and converted into MatLab format for single contraction analysis. Each participant performed four submaximal contractions, with one representative trial chosen for each contraction intensity according to the smoothness and stability of the force trace. Monopolar HDsEMG signals from every contraction were decomposed independently. Signals were band-pass filtered (20-500 Hz, Butterworth, 4^th^ order) and offline decomposed into motor unit pulse trains (MUPTs) by a convolutive kernel compensation algorithm implemented in DEMUSE (Holobar et al., 2014). Following automated decomposition, a trained investigator manually inspected all MU spike trains to verify discharge patterns, correct false detections, and remove spurious signals.

### Analysis of submaximal trapezoidal contraction

All analyses in this section were performed on brief 12-second stable plateau trapezoidal contractions targeting 10%, 25%, 40%, and 50% MVC. From the retained MU spike trains, several key neuromuscular parameters were calculated. Recruitment threshold was defined as the force level (% MVC) at which each MU was first activated during contraction, and derecruitment threshold as the force level at which firing stopped during relaxation. Motor unit discharge rate (MUDR) was assessed as the frequency of motor unit potential (MUP) occurrences within a MUPT, expressed as pulses per second (pps). Discharge variability (CoVISI) was quantified as the coefficient of variation of the interspike interval (ISI) distribution and expressed as a percentage.

### Analysis of triangular ramp contraction

Triangular ramp trials were analysed to characterise dynamic MU firing behaviour across changing force levels. Instantaneous discharge rate was calculated as the inverse of the interspike interval and subsequently smoothed using support vector regression (SVR) with a Gaussian kernel, following the approach previously described (Škarabot et al., 2026). To minimise edge effects during smoothing, the first and last five discharge events within each spike train were assigned weighting coefficients five times greater than those applied to all intermediate discharges. Peak discharge rate determined as the maximum value of the fully smoothed discharge-rate trajectory for each MU. Persistent inward current (PIC) contribution to motoneuron firing was estimated using the paired motor unit technique, whereby onset–offset firing (ΔF) was quantified from matched MU pairs. Each pair consisted of a test MU and a corresponding lower-threshold control MU. Valid MU pairs were required to satisfy the following criteria: (i) rate-to-rate correlation coefficient r^2^ ≥ (ii) test units recruited ≥1 s after control units, and (iii) modulation of the control unit firing rate >0.5 pps. For test MUs paired with multiple valid control MUs, individual ΔF values were averaged to obtain a single representative ΔF estimate for that test

### Analysis of sustained fatiguing contraction

For the sustained 30% MVC fatiguing contraction, MUs were initially decomposed from the first 30s of the contraction to obtain MU filters. These filters were subsequently reapplied throughout the remaining contraction using consecutive 15s overlapping segments, allowing MU activity to be tracked across the progression of fatigue. The transient ramp-up and ramp-down phases were excluded from analysis, and only the stable force plateau phase was retained. Four time points were defined: contraction onset, beginning of the stable phase, end of the stable phase, and contraction offset. The stable contraction period was subsequently divided into five equal-duration phases (P1-5) representing the progression of fatigue from task onset task failure. All phase-wise analyses were performed within these stable contraction intervals, including MUDR and discharge variability (CoVISI). All valid motor units identified within each phase were included in the corresponding phase-specific analyses.

### Blood sampling and biochemical analysis

Blood samples were centrifuged at 3200 rpm for 20 min at 4□. Serum was aliquoted and stored at□−80□ until analysis. Serum caffeine and serotonin concentrations were quantified using competitive ELISA kits (Caffeine ELISA Kit E4558, BioVision, CA, USA; Serotonin ELISA Kit ab133053, Abcam, UK) according to the manufacturers’ instructions. For serotonin, serum samples were diluted 1:16 prior to assay. For caffeine analysis, serum samples underwent manufacturer-recommended pretreatment before dilution. The final 1:1600 dilution factor was determined empirically using pooled serum samples following dilution-linearity and spike-recovery assessments. Analyte concentrations were interpolated from standard curves generated using percentage maximal binding (%B/B0), corrected for dilution and expressed as ng/mL. Inter-plate variation was corrected using internal quality-control samples. Data were logarithmically transformed where necessary to meet normality assumptions.

### Ultrasound

#### Statistics

All statistical processing and data management were implemented within RStudio (Version 1.3.959; R Foundation for Statistical Computing, Vienna, Austria). Statistical significance was set at p = 0.05. Linear mixed-effects models (LMMs) were fitted using restricted maximum likelihood (REML) with the lme4 package (Bates et al., 2015). Type III analyses of variance (ANOVA) with Satterthwaite’s degrees-of-freedom approximation were performed using the lmerTest package to evaluate main effects and interaction terms (Kuznetsova et al., 2017). Model assumptions were assessed visually using quantile-quantile plots and residual diagnostics.

For circulating blood biomarkers, LMMs were performed with Intervention (PLA, CAFF), TimePoint (B0, P2, P3), and their interaction as fixed effects, with participant included as a random intercept. For short trapezoidal contractions, LMMs were fitted with Intervention (PLA, CAFF) and Contraction Level (10%, 25%, 40% and 50% MVC), and their interaction as fixed effects. For triangular ramp contractions, LMMs were used to assess intervention effects on ramp-derived MU variables, including peak discharge rate and ΔF. For analyses of short trapezoidal and triangular ramp contractions, MUs were not tracked across contraction levels. Therefore, participant was included as the sole random effect, while recruitment threshold was included as a covariate to account for differences in MU recruitment characteristics.

For sustained fatiguing contractions, both categorical phase-based and polynomial mixed-effects models were initially evaluated. Model fit was assessed using Akaike Information Criterion (AIC) and Bayesian Information Criterion (BIC). Polynomial models provided a more parsimonious description of fatigue-related trajectories for MU discharge variables and were therefore retained for final analyses. Specifically, fatigue-related trajectories in MU discharge rate and discharge variability were analysed using polynomial mixed-effects models. Fatigue phase was treated as a continuous variable from P1 to P5 and modelled using a second-order polynomial term. Models included Intervention, the polynomial phase term, and their interaction as fixed effects. Recruitment threshold relative to MVC was included as a covariate for MU discharge outcomes. As individual MUs were measured repeatedly across fatigue phases, random intercepts were included for participant and MU nested within participant.

Estimated marginal means (EMMs) and standard errors (SEs) were derived from the final fitted mixed-effects models using the emmeans package (Lenth & Piaskowski, 2017). Where significant main effects or interactions were detected, post hoc pairwise comparisons were performed using Tukey-adjusted multiple comparisons.

Exploratory correlation analyses were conducted to determine whether inter-individual variability in caffeine exposure was associated with the magnitude of caffeine-induced neuromuscular responses. Caffeine-induced changes (Δ) were calculated as the difference between caffeine and placebo conditions at the corresponding post-ingestion time points. Motor unit outcomes were averaged within each participant and condition prior to calculating Δvalues. Associations between changes in circulating caffeine concentration and changes in motor unit outcomes were assessed using Spearman’s rank correlation coefficients.

## Results

Seventeen participants completed the study. Participant characteristics are presented in Table 1.

**Table 1.** Participant characteristics. Values are presented as mean ± SD.

| Characteristic | Value |
| --- | --- |
| Participants, n | 17 |
| Age (years) | 22.6 ± 4.4 |
| BMI (kg/m<sup>2</sup>) | 25.6 ± 3.2 |
| Caffeine dose (mg) | 240.8 ± 39.3 |
| VL CSA (cm<sup>2</sup>) | 31.4 ± 6.1 |

Abbreviations: BMI, body mass index; VL, vastus lateralis; CSA, cross-sectional area.

### Blood results

Circulating caffeine concentration demonstrated significant main effects of Intervention (F (1,72) = 321.10, p<0.001) and TimePoint (F (2,72) = 66.50, p<0.001), as well as a significant Intervention × TimePoint interaction (F (2,72) = 90.10, p<0.001) (Fig 2A). Pairwise comparisons revealed no difference between PLA and CAFF at baseline (mean difference = 193 ng/mL, p=0.584). In contrast, circulating caffeine concentration was markedly greater under CAFF than PLA at P2 (mean difference = 5117 ng/mL, p<0.001) and P3 (mean difference = 5964 ng/mL, p<0.001). Within the placebo condition, caffeine concentration remained unchanged across time (all p>0.49). Within the caffeine condition, caffeine concentration increased substantially from baseline to P2 (mean difference = 4927 ng/mL, p<0.001) and remained elevated at P3 (mean difference = 5742 ng/mL¹, p<0.001), with P3 values also exceeding P2 (mean difference = 815 ng/mL, p=0.045).

**Figure 1.**
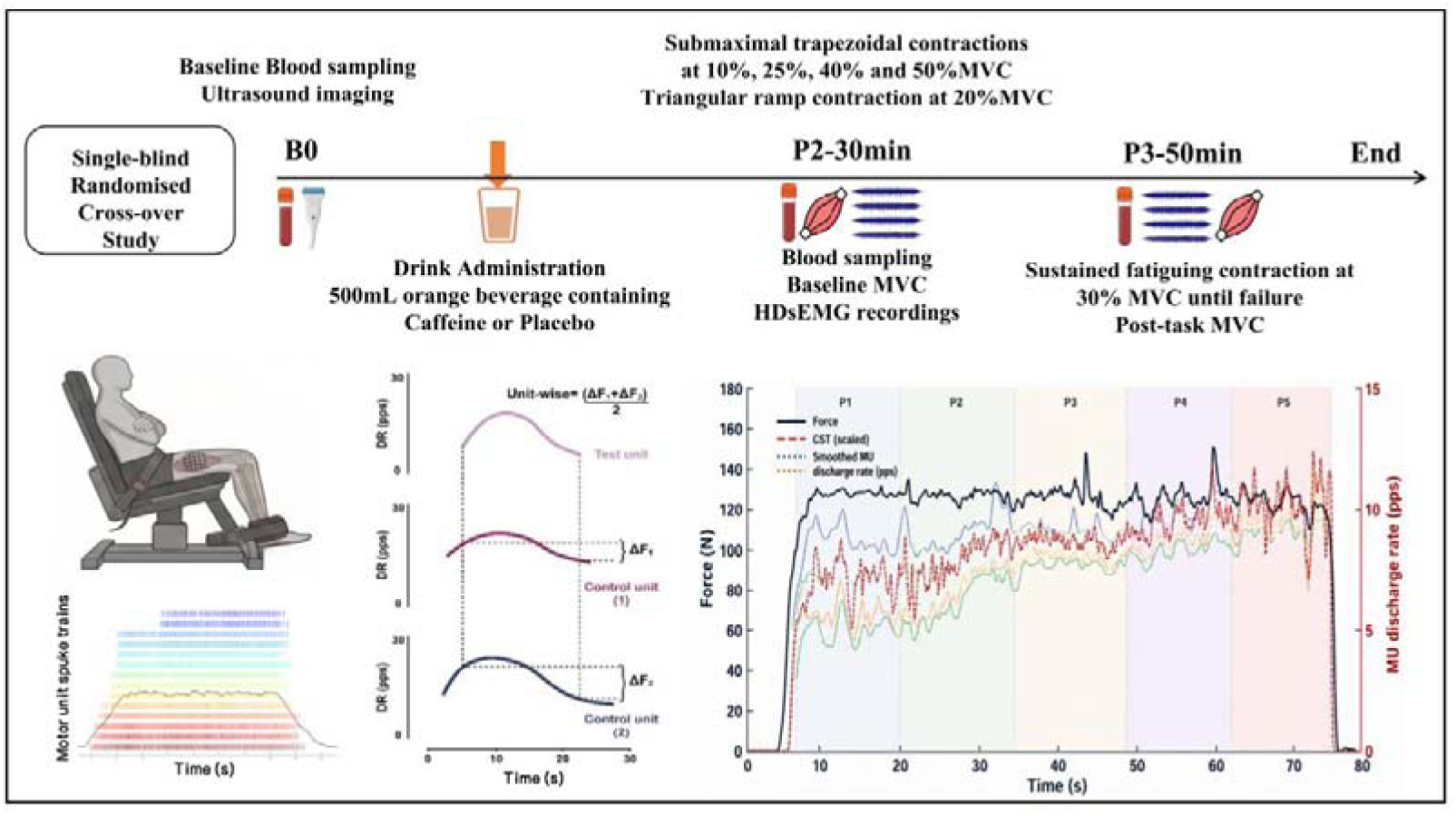
Experimental protocol and overview of motor unit analyses. Participants completed two experimental sessions in a randomised, single-blind, placebo-controlled crossover design. At baseline (B0), blood samples were collected and ultrasound imaging was performed before ingestion of either a caffeine-containing or placebo orange beverage. Thirty minutes after ingestion (P2), a second blood sample was obtained, followed by high-density surface electromyography (HDsEMG) recordings during graded trapezoidal contractions (10%, 25%, 40%, and 50% MVC) and a triangular ramp contraction (20% MVC) for motor unit discharge and persistent inward current (ΔF) analyses. Fifty minutes after ingestion (P3), a third blood sample was collected prior to a sustained isometric contraction performed at 30% MVC until task failure, during which motor unit discharge behaviour was analysed. Representative examples of the experimental setup, motor unit decomposition, ΔF estimation, and phase-based analysis of the sustained fatiguing contraction are shown in the lower panels. MVC, maximal voluntary contraction.

**Figure 2.**
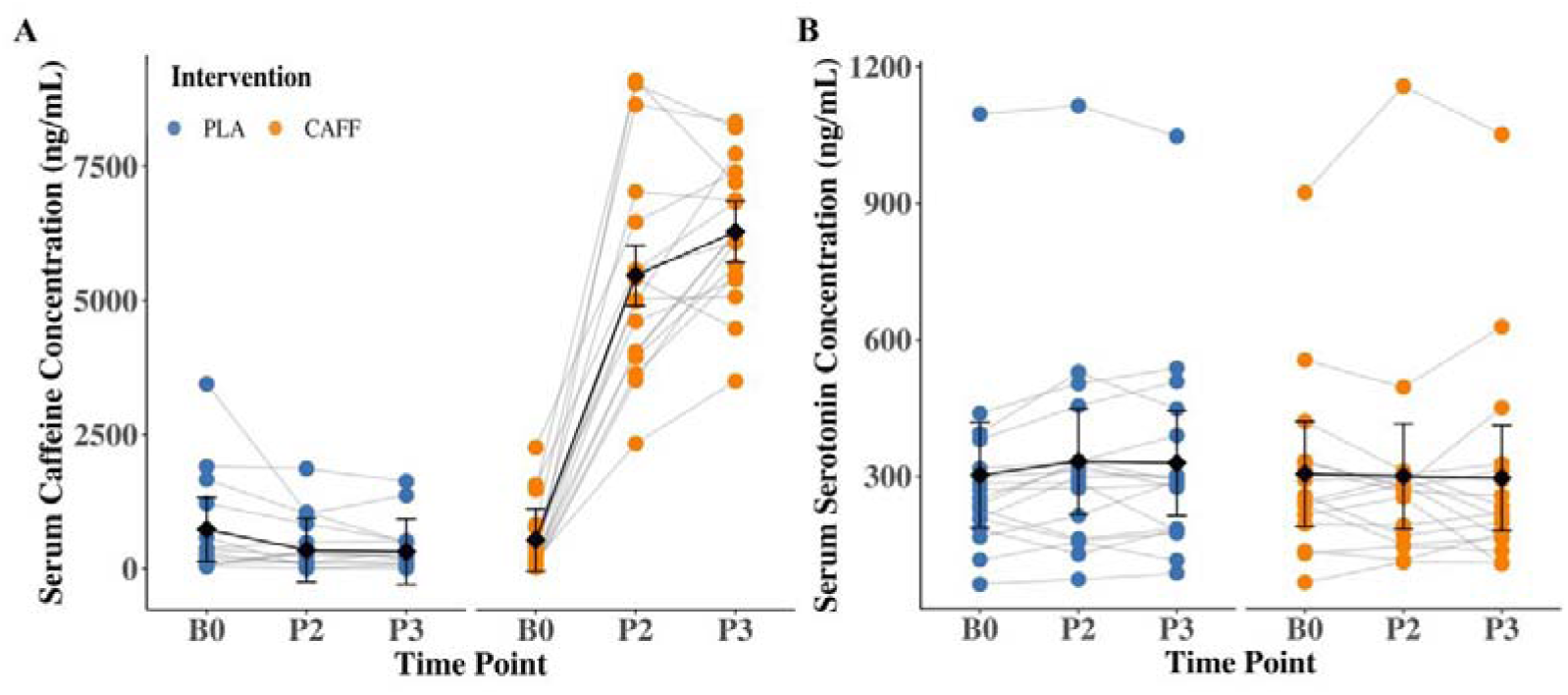
Circulating serum caffeine and serotonin concentrations over time in placebo (PLA) and caffeine (CAFF) conditions. Individual participant data are displayed as grey connected lines with overlaid points, illustrating within-subject temporal trajectories. Estimated marginal means (EMMs) derived from the mixed-effects models are shown as black diamond markers, with vertical error bars representing 95% confidence intervals. Solid black lines indicate model-estimated temporal trends across time points.

Circulating serotonin concentration did not demonstrate a significant main effect of Intervention (F (1,80) = 3.16, p=0.079), TimePoint (F (2,80) = 0.40, p=0.669), or Intervention × TimePoint interaction (F (2,80) = 1.01, p=0.369) (Fig 2B).

### Functional Performance

Baseline maximal voluntary contraction (MVC) was comparable between caffeine and placebo conditions (CAFF: 521.68 ± 145.71 N; PLA: 502.24 ± 164.92 N) (Fig 3A). MVC was lower at the post-task assessment in both conditions (main effect of Time: F(1,48) = 144.43, p < 0.001), with no significant main effect of Intervention (F(1,48) = 3.21, p = 0.079) or Intervention × Time interaction (F(1,48) = 0.036, p = 0.850). Thus, the baseline-to-post reduction in MVC did not differ between caffeine and placebo conditions.

**Figure 3.**
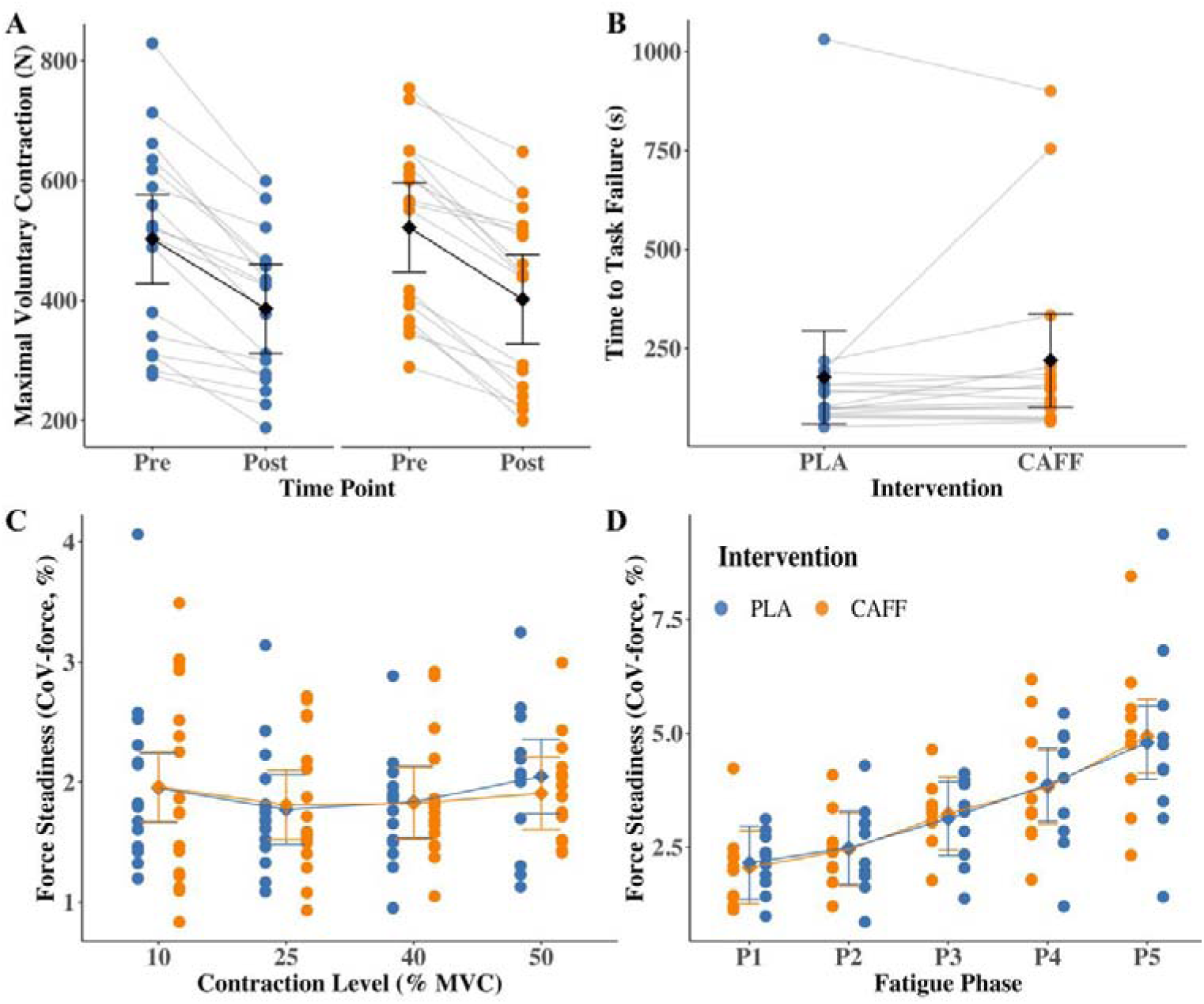
Physical function-related measurements in placebo (PLA) and caffeine (CAFF) conditions. Individual participant data are displayed in coloured points and grey lines indicate individual responses. Estimated marginal means (EMMs) derived from the mixed-effects models are shown as diamond markers, with vertical error bars representing 95% confidence intervals. Solid black lines indicate model-estimated temporal trends across time points.

There was no signficant difference in Time to task failure between the caffeine and placebo conditions (219.96 ± 239.23 vs. 177.15 ± 225.13 s, p = 0.237) (Fig 3B).

Force steadiness, quantified as the coefficient of variation of force output (CoV-force), was analysed separately for submaximal trapezoidal contractions and sustained fatiguing contractions. During submaximal trapezoidal contractions, no significant main effect of Intervention was observed (F (1,102.09) = 0.13, p=0.714), nor was there a significant main effect of Force Level (F (3,102.41) =1.63, p=0.188) or an Intervention × Level interaction (F (3,101.98) = 0.28, p=0.838) (Fig 3C), indicating that caffeine ingestion did not influence force steadiness across different contraction intensities under steady-state conditions. In contrast, during sustained fatiguing contractions, a significant main effect of Fatigue Phase was observed (F (2,85) = 60.08, p<0.001), indicating a progressive increase in force fluctuations throughout the contraction. Neither the main effect of Intervention (F (1,85) = 0.03, p=0.859) nor the Intervention × Phase interaction (F (2,85) = 0.07, p=0.931) was statistically significant (Fig 3D). CoV-force increased progressively from the initial to the final phase under both conditions, reflecting the expected decline in force steadiness with fatigue development.

### Submaximal contraction

Valid MU decompositions were obtained from all 17 participants at 10% and 25% MVC, 15 participants at 40% MVC, and 13 participants at 50% MVC. Across these contraction levels, 294/293, 254/244, 182/168, and 129/140 MUs were identified in the placebo/caffeine conditions, respectively, corresponding to mean yields of 17 ± 12/17 ± 12, 15 ± 12/14 ± 13, 12 ± 11/11 ± 10, and 10 ± 8/9 ± 8 MUs per participant.

### Motor unit discharge rate and discharge variability

Significant main effects of Intervention (F (1, 1680) = 32.75, p < 0.001) and Contraction Level (F (3, 1680) = 857, p < 0.001) were observed in MU discharge rate (MUDR) (Fig 4A). The Intervention × Level interaction was not significant (F (3,1680) = 0.76, p=0.518), with MUDR being higher in the caffeine condition compared with placebo across all contraction intensities. Estimated marginal means were consistently greater under caffeine than placebo at 10% MVC (CAFF vs. PLA; 6.94 ± 0.39 vs. 6.64 ± 0.39 pps, p = 0.010), 25% MVC (9.55 ± 0.39 vs. 9.19 ± 0.39 pps, p = 0.004), 40% MVC (12.18 ± 0.39 vs. 11.60 ± 0.39 pps, p < 0.001), and 50% MVC (14.77 ± 0.40 vs. 14.38 ± 0.40 pps, p = 0.022).

**Figure 4.**
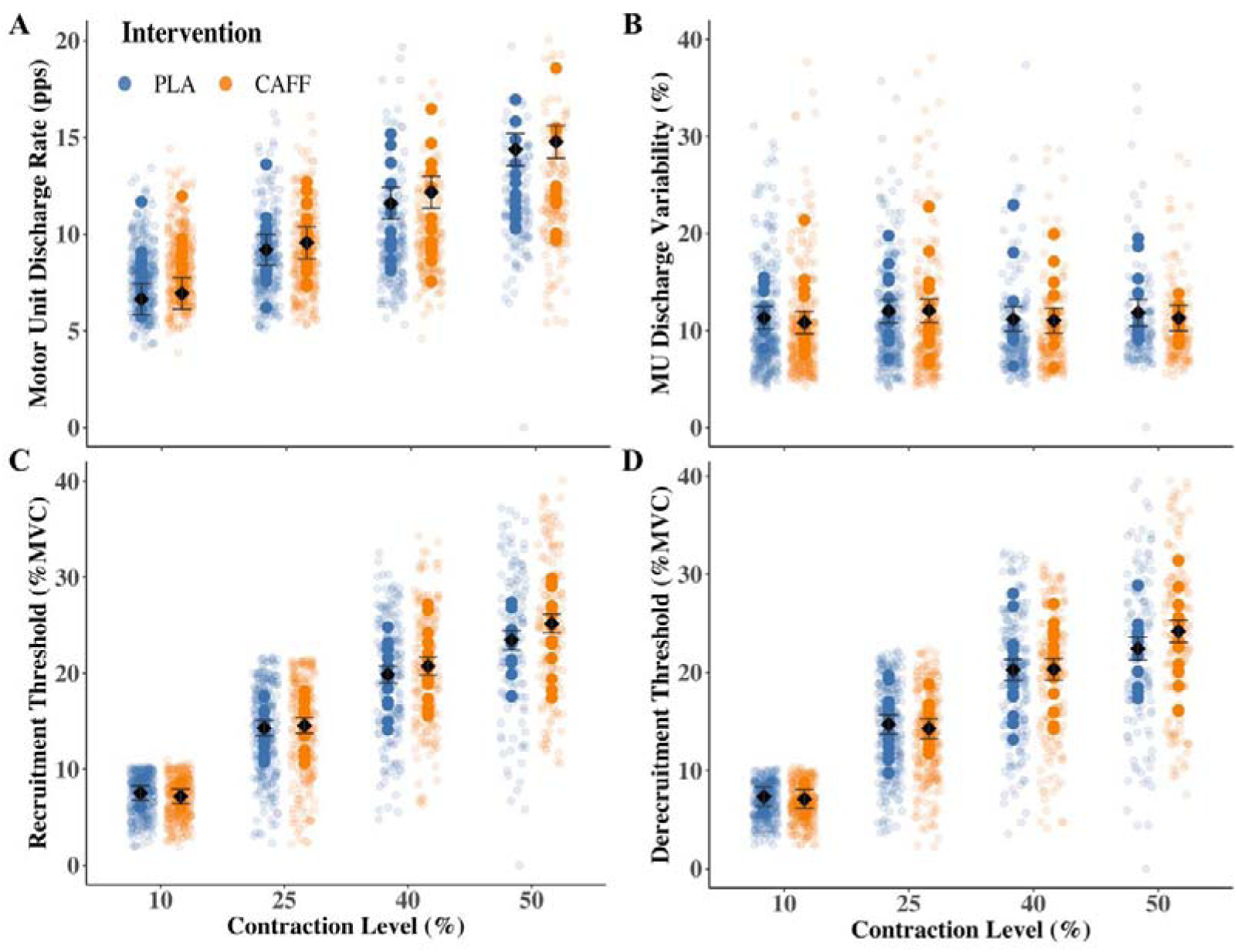
Differences in motor unit discharge properties between placebo (PLA) and caffeine (CAFF) conditions during short trapezoidal contractions at 10%, 25%, 40% and 50% of maximal voluntary contraction (MVC). (A) Motor unit (MU) discharge rate (mean discharge rate, in pulses per second, pps). (B) MU discharge variability (coefficient of variation of inter-spike intervals, CoVISI; in %). (C) Motor unit recruitment threshold expressed relative to MVC (%MVC). (D) Motor unit derecruitment threshold expressed relative to MVC (%MVC). Individual MU observations are displayed as semi-transparent jittered points, while participant-level means are shown as larger coloured circles. The horizontal black diamonds indicate estimated marginal means derived from linear mixed-effects models, with associated 95% confidence intervals represented by vertical error bars. Placebo and caffeine conditions are displayed in blue and orange separately at each contraction level.

Similarly, MU discharge variability demonstrated significant main effects of Intervention (F (1,1684) = 5.77, p=0.016) and Contraction Level (F (3,1683) = 84.60, p<0.001) (Fig 4B). The Intervention × Level interaction was not significant (F (3,1681) = 1.22, p=0.302), with discharge variability being lower in the caffeine condition compared with placebo across contraction levels. Estimated marginal means were lower under caffeine than placebo at all contraction levels, including 10% MVC (14.48 ± 0.56 vs. 14.77 ± 0.56%, p = 0.467), 25% MVC (12.04 ± 0.55 vs. 12.12 ± 0.54%, p = 0.854), 40% MVC (7.96 ± 0.61 vs. 8.53 ± 0.60%, p = 0.267), and 50% MVC (6.03 ± 0.67 vs. 7.45 ± 0.66%, p = 0.016).

Exploratory Spearman correlation analyses showed that changes in circulating caffeine concentration were not significantly associated with caffeine-induced changes in MU discharge properties. No significant correlations were observed between Δcaffeine concentration and Δmean MUDR (ρ = 0.19, p = 0.529) or ΔMU discharge variability (ρ = −0.40, p = 0.176).

### Recruitment/Derecruitment Threshold (%MVC)

For MU recruitment threshold expressed relative to MVC, significant main effects of Intervention (F (1,1691) = 6.43, p=0.011) and Contraction Level (F (3,1692) = 911.11, p<0.001) were observed (Fig 4C). A significant Intervention × Level interaction was also detected (F (3, 1686) = 3.11, p = 0.026), indicating that the effect of caffeine on recruitment threshold varied across contraction levels. Estimated marginal means revealed no differences between caffeine and placebo at 10% MVC (7.17 ± 0.38 vs. 7.52 ± 0.38 %MVC, p=0.391), 25% MVC (14.56 ± 0.40 vs. 14.30 ± 0.40 %MVC, p=0.556), or 40% MVC (20.76 ± 0.46 vs. 19.88 ± 0.45 %MVC, p=0.092). However, recruitment thresholds were significantly greater under caffeine at 50% MVC (25.18 ± 0.49 vs. 23.45 ± 0.51 %MVC, p=0.004).

For motor unit derecruitment threshold expressed relative to MVC, a significant main effect of Contraction Level was observed (F (3,1688) = 882.42, p<0.001) (Fig 4D). Although the main effect of Intervention was not significant (F (1,1687) = 1.44, p=0.230), a significant Intervention × Level interaction was detected (F (3,1684) = 3.41, p=0.017), indicating that intervention effects differed across contraction intensities. Estimated marginal means showed no differences between caffeine and placebo at 10% MVC (7.09 ± 0.48 vs. 7.33 ± 0.48 %MVC, p=0.543), 25% MVC (14.29 ± 0.50 vs. 14.70 ± 0.49 %MVC, p=0.340), or 40% MVC (20.32 ± 0.54 vs. 20.26 ± 0.53 %MVC, p=0.899). In contrast, derecruitment thresholds were significantly greater under caffeine at 50% MVC (24.19 ± 0.57 vs. 22.42 ± 0.58 %MVC, p=0.003).

### ΔF and Peak Discharge Rate

Valid MU decompositions were obtained from 12 participants during the 20% MVC ramp contraction. A total of 207 and 217 MUs were identified in the placebo and caffeine conditions, respectively, corresponding to 17 ± 10 and 18 ± 12 MUs per participant.

For Δ**F**, there was no significant main effect of intervention between the caffeine and placebo conditions (F (1, 326) = 3.01, 2.15 ± 0.22 vs.2.36 ± 0.22, p = 0.083) (Fig 5A).

**Figure 5.**
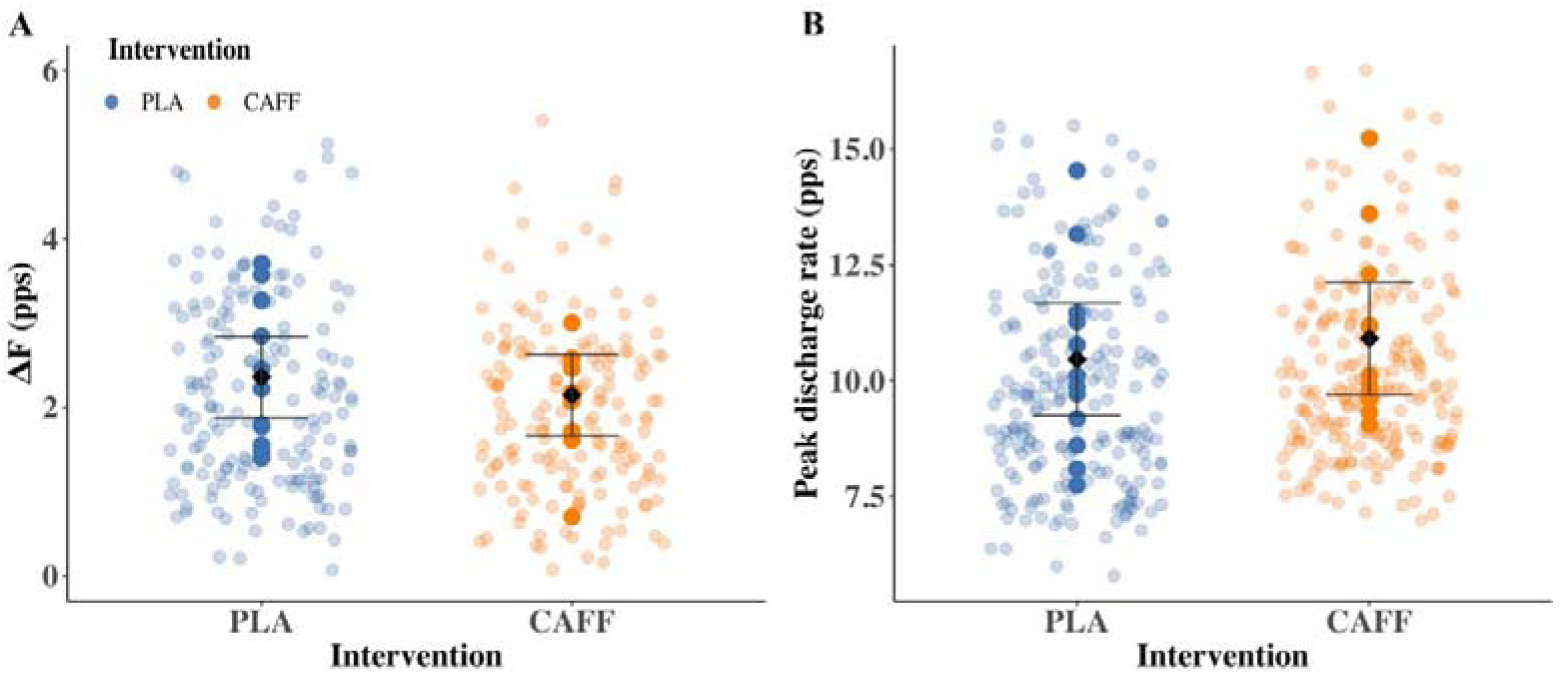
Differences in motor unit discharge hysteresis (ΔF) and peak discharge rate between placebo (PLA) and caffeine (CAFF) conditions during triangular ramp contractions at 20% of maximal voluntary contraction (MVC). Individual motor unit (MU) observations are displayed as semi-transparent jittered points, whereas participant-level means are shown as larger coloured circles. The horizontal black diamonds represent estimated marginal means derived from linear mixed-effects models, with 95% confidence intervals indicated by vertical error bars. Placebo and caffeine conditions are displayed in blue and orange separately at the group level.

Peak discharge rate was significantly higher in the caffeine condition than in the placebo condition. Estimated marginal means were 10.90 ± 0.56 pps for CAFF and 10.34 ± 0.56 pps for PLA. The mixed-effects model revealed a significant main effect of intervention (F (1,410) = 17.79, p < 0.001) (Fig 5B).

### Sustained fatiguing contraction

Valid MU decompositions were obtained from 10 participants during the sustained fatiguing contraction at 30% MVC, with a total of 87 and 79 MUs identified in the placebo and caffeine conditions, respectively, with an average of 9 ± 4 and 8 ± 5 MUs available per participant per fatigue phase.

A polynomial mixed-effects model revealed a significant effect of Fatigue Phase on MUDR (F (2, 696) = 52.47, p < 0.001), indicating that MUDR changed throughout the fatiguing contraction (Fig 6A). No significant effect of Intervention was observed (F (1, 697) = 2.17, p = 0.141). However, a significant Intervention × Phase interaction was detected (F (2, 695) = 5.02, p = 0.007), indicating that the temporal trajectory of MUDR differed between conditions. Estimated marginal means showed no differences between caffeine and placebo during P1 (8.69 ± 0.38 vs. 8.47 ± 0.37 pps, p=0.247) or P2 (8.28 ± 0.36 vs. 8.38 ± 0.36 pps, p=0.429). In contrast, MUDR was significantly lower under caffeine during P3 (8.24 ± 0.36 vs. 8.56 ± 0.36 pps, p=0.015), P4 (8.56 ± 0.36 vs. 9.02 ± 0.36 pps, p<0.001), and P5 (9.24 ± 0.37 vs. 9.76 ± 0.37 pps, p=0.003). Across both conditions, MUDR initially declined from P1 to P2 before progressively increasing toward P5, reaching its highest values during the final phase of the contraction.

**Figure 6.**
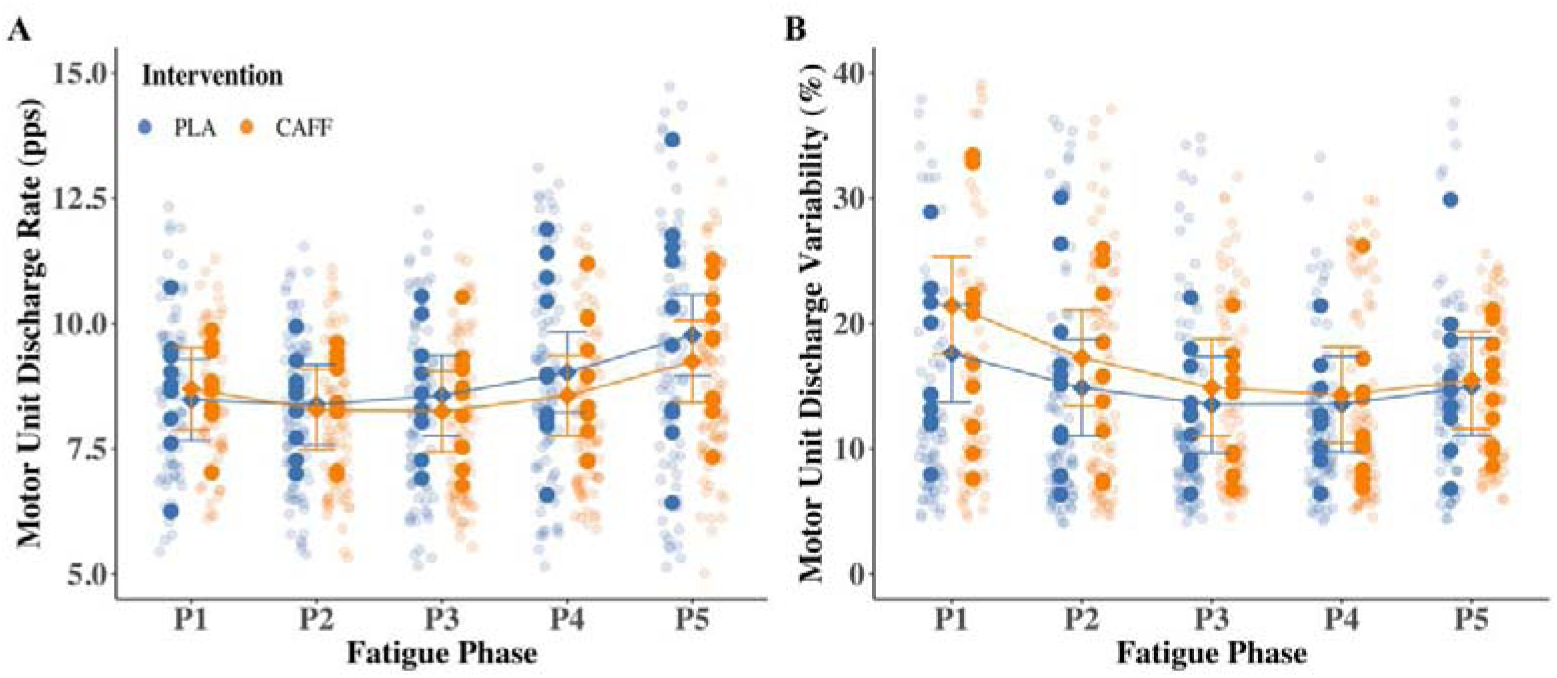
Motor unit discharge rate and discharge variability across fatigue phases during sustained isometric contractions. Panels illustrate fatigue-related changes in motor unit discharge properties across five equally divided phases (P1–P5) of a sustained contraction performed at 30% MVC. Data are shown for placebo (PLA) and caffeine (CAFF) conditions. Individual motor unit (MU) observations are displayed as semi-transparent jittered points. Subject-level means are shown as larger coloured circles. Estimated marginal means (EMMs) derived from polynomial mixed-effects models are indicated by diamond markers, with vertical bars representing 95% confidence intervals. Continuous fatigue trajectories were modelled using second-order polynomial mixed-effects models and are shown as condition-specific trend lines connecting EMMs across phases.

For MU discharge variability, significant main effects of Intervention (F (1,696) = 8.36, p=0.004) and Fatigue Phase (F (2,696) = 48.32, p<0.001) were observed (Fig 6B). A significant Intervention × Phase interaction was also detected (F (2,694) = 4.55, p=0.011), indicating that temporal changes in discharge variability differed between conditions. Estimated marginal means revealed greater discharge variability under caffeine during P1 (21.4 ± 1.79 vs. 17.6 ± 1.77%, p<0.001), P2 (17.3 ± 1.72 vs. 14.9 ± 1.71%, p<0.001), and P3 (14.9 ± 1.73 vs. 13.5 ± 1.72%, p=0.029). No between-condition differences were observed during P4 (14.3 ± 1.72 vs. 13.5 ± 1.70%, p=0.181) or P5 (15.5 ± 1.77 vs. 14.9 ± 1.76%, p=0.530). Across both interventions, discharge variability declined from P1 to the middle phases of the contraction before increasing again toward task failure, demonstrating a U-shaped fatigue trajectory.

## Discussion

This study characterised the effects of acute caffeine ingestion on circulating serum caffeine levels and neuromuscular function during submaximal and fatiguing contractions. Caffeine did not alter maximal force production, fatigue resistance, or force steadiness, but increased MU discharge rate during graded contractions and modified the trajectories of MU discharge rate and variability during sustained fatigue. ΔF remained unchanged, providing no evidence for a measurable alteration in PIC-related intrinsic motoneuron excitability under the present experimental conditions. Collectively, these findings indicate that caffeine alters MU behaviour in a task-dependent manner without producing corresponding improvements in whole-task performance.

Although acute caffeine ingestion is widely recognised to elicit ergogenic benefits across diverse exercise modalities (Grgic et al., 2018), the present study observed no improvements in maximal force capacity, endurance performance, or force steadiness during either trapezoidal or sustained fatiguing isometric contractions. Importantly, serum caffeine concentrations confirmed robust systemic absorption, indicating that the absence of performance effects was not attributable to insufficient bioavailability. These findings are broadly consistent with previous reports (Pickering & Grgic, 2019), reporting limited or inconsistent ergogenic effects of caffeine during isometric strength tasks, yet contrast with the extensive literature documenting consistent caffeine-induced enhancements in exercise performance.

The existing literature demonstrates substantial heterogeneity in caffeine’s ergogenic effects across muscle groups and exercise modalities (Grgic et al., 2020), with more pronounced benefits often reported in upper-limb compared with lower-limb tasks (Grgic et al., 2018), and variable effects even within lower-limb extensor and flexor muscles (Warren et al., 2009). Similarly, evidence regarding endurance performance remains mixed, with some studies reporting enhanced time-to-exhaustion and better performance (Kalmar & Cafarelli, 1999), whereas others show minimal or no effect, particularly in untrained populations (Jodra et al., 2020; Khodadadi et al., 2025). Beyond changes in force output, caffeine-related effects on the ability to maintain steady force production remain comparatively unclear, with limited evidence for consistent modulation of force variability during sustained contractions. Given the absence of overt performance effects despite serum-confirmed caffeine uptake, the mechanistic focus shifts toward the level of motor unit behaviour, which may reveal subclinical neural adaptations not reflected in global force output.

Previous studies have reported variable effects of acute caffeine ingestion on MU behaviour, with changes differing according to the muscle examined and contraction task (Ghazaleh et al., 2024; Lin et al., 2025). In the present study, caffeine increased mean MU discharge rates across all contraction levels and was associated with an overall reduction in MU discharge variability during submaximal contractions. These findings indicate that caffeine modulates MU output in an intensity-dependent and non-uniform manner, influencing both discharge rate and discharge variability depending on functional demand. MU discharge rate represents the net output of the motoneuron pool and is influenced by both descending drive and intrinsic motoneuron properties. Acute caffeine acts primarily as an antagonist of adenosine A_1_ and A_2_A receptors, reducing the inhibitory influence of adenosine on cortical neurons and thereby increasing cortical excitability (Nehlig et al., 1992). This enhancement in cortical excitability is thought to facilitate corticospinal output and augment descending excitatory drive to spinal motoneurons, providing a plausible physiological basis for higher discharge rates observed in the present study. However, the present measurements cannot localize the neural source of this altered MU output.

Previous studies provide mixed but partially convergent evidence regarding caffeine’s effects on MU behaviour. Increases in EMG amplitude have been reported following caffeine ingestion, consistent with enhanced neural activation during voluntary contractions (Amoruso et al., 2026). For example, reductions in discharge variability and improvements in force precision have been reported during isometric contractions of the tibialis anterior (Lin et al., 2025), whereas studies in the vastus lateralis have reported minimal or no changes in MU discharge behaviour or EMG–force relationships (Kalmar & Cafarelli, 1999), highlighting muscle-dependent variability in caffeine responsiveness. Consistent with this, caffeine ingestion has been reported to produce no detectable changes in EMG activity in lower-limb muscles (Ghazaleh et al., 2024), further emphasising that caffeine-induced neural adaptations are not uniformly expressed across muscles or contraction conditions.

Beyond discharge rate and its variability, recruitment behaviour also shows evidence of caffeine-induced modulation at higher force levels. We observed an increase in recruitment threshold at higher force levels under caffeine ingestion, indicating a selective modulation of higher-threshold MU recruitment that emerges under elevated contraction demands. This effect aligns with previously reported non-linear recruitment behaviour under caffeine (Mackay et al., 2023) and is consistent with the broader evidence of muscle- and intensity-dependent modulation of recruitment thresholds (Nishikawa et al., 2024). Collectively, these results suggest that acute caffeine primarily influences MU behaviour by increasing discharge rate while improving discharge regularity, particularly under conditions of greater neural demands (Walton et al., 2002).

To determine whether inter-individual variability in caffeine exposure influenced MU responses, exploratory correlations were performed between caffeine-induced changes in circulating caffeine concentration and changes in MU discharge properties. No significant associations were observed between changes in caffeine concentration and changes in either mean MU discharge rate or discharge variability. These findings suggest that circulating caffeine concentration alone does not account for the observed inter-individual variability in MU responses.

We found no difference in ΔF between caffeine and placebo conditions at 20% MVC, indicating acute caffeine ingestion did not alter intrinsic motoneuron excitability under the present experimental conditions. Circulating serotonin concentrations likewise remained unchanged, providing no evidence for a systemic serotonergic response accompanying the unchanged ΔF. However, circulating serotonin does not directly reflect serotonergic signalling within the central nervous system or serotonergic input to spinal motoneurons; therefore, the absence of a serum response should not be interpreted as direct evidence that central serotonergic modulation was unchanged. Evidence regarding caffeine-related modulation of PICs remains inconsistent across the literature. Some studies have reported increased indices of motoneuron excitability following caffeine ingestion, including elevated ΔF and enhanced self-sustained firing, suggesting a potential facilitation of persistent motoneuron activity under certain conditions (Popesco et al., 2026; Walton et al., 2002). However, other investigations have reported no changes in PIC amplitudes or ΔF following caffeine ingestion across different contraction intensities (Kirk et al., 2019; Mackay et al., 2023), highlighting substantial variability in reported effects across contexts. Taken together, the absence of changes in ΔF provides no evidence that acute caffeine ingestion altered intrinsic motoneuron excitability under the present experimental conditions, while the unchanged circulating serotonin concentrations indicate only the absence of a detectable systemic serotonergic response. This finding suggests that the increases in MU discharge observed following caffeine ingestion are unlikely to be explained by altered intrinsic motoneuron excitability, prompting further consideration of how caffeine influences MU behaviour during progressive fatigue.

In addition to the intensity-dependent MU modulation observed during submaximal trapezoidal contractions, caffeine-induced effects were also evident in the sustained fatiguing protocol. Net excitation to the motoneuron pool is known to increase during fatigue to compensate for reductions in MU discharge and to maintain target force output (Adam & De Luca, 2005; Contessa et al., 2016; McManus et al., 2015; Vila-Cha et al., 2012). Consistent with this framework, increases in MU discharge rate have been reported during fatiguing contractions in multiple muscles, reflecting progressive increases in central drive as task demands accumulate (Gomes et al., 2025). However, MU discharge behaviour does not increase linearly throughout sustained contraction. Previous studies have reported progressive declines in mean MU discharge rate as fatigue accumulates (Conwit et al., 2000), while subsequent work demonstrated that cumulative fatigue can alter the initial discharge state of later contractions, suggesting that the motoneuron pool undergoes dynamic reorganisation across repeated efforts. Extending this concept, recent studies have identified a biphasic pattern of MU discharge modulation during sustained contractions, characterised by an initial decline followed by a progressive increase in discharge rate as task failure approaches (Martinez-Valdes et al., 2020, 2026; Valenčič et al., 2024). Consistent with this finding, polynomial modelling in the present study identified a similar biphasic trajectory, indicating that MU output is dynamically regulated rather than progressively declining throughout fatigue development.

MU discharge variability has also been shown to change with fatigue, although the direction and magnitude of these effects are not uniform across studies (Contessa et al., 2009, 2018). Given that endogenous MU discharge patterns are dynamically reconfigured across fatigue phases, caffeine-related effects observed in the present study should be interpreted within the context of this dynamically evolving neuromuscular state. In the present study, caffeine modified the temporal trajectories of both MU discharge rate and discharge variability during the sustained contraction, indicating that its influence on MU output changes across fatigue progression. Together with the alterations observed during graded contractions, these findings suggest that caffeine does not exert a uniform effect on MU behaviour across contraction conditions.

## Strengths and Limitations

This study provides a multi-level characterisation of the neuromuscular effects of acute caffeine ingestion by integrating circulating caffeine levels, MU discharge behaviour, and estimates of motoneuron intrinsic excitability across both submaximal and fatiguing isometric contractions. The inclusion of both trapezoidal and sustained fatiguing contraction protocols further allowed evaluation of caffeine effects across distinct neuromuscular states, providing insight into intensity- and fatigue-dependent modulation of MU behaviour. However, several limitations should be considered when interpreting the present findings. (1) While serum caffeine concentrations confirmed systemic absorption, inter-individual variability in adenosine receptor sensitivity may have contributed to variability in neuromuscular responses, which was not assessed in the present analysis. Similarly, serum serotonin concentrations are not a direct measure of bioavailable serotonin influencing MN behaviour. (2) Estimates of persistent inward currents were derived from ΔF analysis during low-intensity contractions, which, although widely used, provides only an indirect index of intrinsic motoneuron excitability and may not fully capture caffeine-related changes occurring under different conditions. (3) The present findings are based exclusively on isometric contractions of vastus lateralis in young healthy individuals, and therefore may not generalise to dynamic movements, different muscle groups, or populations with altered neuromuscular function.

## Conclusions

Acute caffeine ingestion did not enhance maximal force production, force steadiness, or fatigue resistance during isometric knee-extensor contractions, despite producing clear changes in MU behaviour. Caffeine increased MU discharge rate during graded contractions, altered discharge variability, and modified the temporal trajectory of MU firing during sustained fatigue. These changes occurred without a detectable change in ΔF, providing no evidence for altered PIC-related intrinsic motoneuron excitability under the present experimental conditions. Overall, caffeine modulates motor unit output in an intensity- and fatigue-dependent manner, but these neural changes do not translate into improved whole-task performance.

## Acknowledgments

We thank Mrs Amanda Gates for her valuable assistance with blood collection. We also thank all of the participants for their time and participation.

## Author Contributions

All authors contributed to the conception and design of the study. YG, TI, EJ and VS contributed to the data acquisition. YG and MP analysed the data and drafted the manuscript. BEP and PJA provided comments. All authors have approved the final version of the submitted manuscript for publication and are accountable for all aspects of the work. All persons designated as authors qualify for authorship, and all those who qualify for authorship are listed.

## Data availability statement

The datasets generated and analysed during the current study are available from the corresponding author upon reasonable request.

